# A First-in-class, Highly Selective and Cell-active Allosteric Inhibitor of Protein Arginine Methyltransferase 6 (PRMT6)

**DOI:** 10.1101/2020.12.04.412569

**Authors:** Yudao Shen, Fengling Li, Magdalena M. Szewczyk, Levon Halebelian, Irene Chau, Mohammad S. Eram, Carlo Dela Seña, Kwang-Su Park, Fanye Meng, He Chen, Hong Zeng, David McLeod, Carlos A. Zepeda-Velázquez, Robert M. Campbell, Mary M. Mader, Brian M. Watson, Matthieu Schapira, Cheryl H. Arrowsmith, Rima Al-Awar, Dalia Barsyte-Lovejoy, H. Ümit Kaniskan, Peter J. Brown, Masoud Vedadi, Jian Jin

## Abstract

PRMT6 catalyzes monomethylation and asymmetric dimethylation of arginine residues in various proteins, plays important roles in biological processes and is associated with multiple cancers. While there are several reported PRMT6 inhibitors, a highly selective PRMT6 inhibitor has not been reported to date. Furthermore, allosteric inhibitors of protein methyltransferases are rare. Here we report the discovery and characterization of a first-in-class, highly selective allosteric inhibitor of PRMT6, SGC6870. SGC6870 is a potent PRMT6 inhibitor (IC_50_ = 77 ± 6 nM) with outstanding selectivity for PRMT6 over a broad panel of other methyltransferases and non-epigenetic targets. Notably, the crystal structure of the PRMT6–SGC6870 complex and kinetic studies revealed SGC6870 binds a unique, induced allosteric pocket. Additionally, SGC6870 engages PRMT6 and potently inhibits its methyltransferase activity in cells. Moreover, SGC6870’s enantiomer, SGC6870N, is inactive against PRMT6 and can be utilized as a negative control. Collectively, SGC6870 is a well-characterized PRMT6 chemical probe and valuable tool for further investigating PRMT6 functions in health and disease.

## INTRODUCTION

Protein arginine methyltransferases (PRMTs) catalyze monomethylation and symmetric or asymmetric dimethylation of arginine residues at histone and/or non-histone substrates.^1, 2^ PRMTs are involved in multiple cellular processes, including regulation of cell cycle, cell pluripotency, estrogen-stimulated transcription, DNA repair, and modulation of pre-mRNA splicing.^3, 4^ Aberrant expression of PRMTs are associated with various types of cancer, such as leukemia, breast, lung, prostate and bladder cancers.^5^ PRMTs are commonly grouped into three types.^1, 2^ Type I PRMTs including PRMT1, PRMT3, PRMT4 (CARM1), PRMT6 and PRMT8 catalyze monomethylation and asymmetric dimethylation of arginine residues. Type II PRMTs including PRMT5 and PRMT9 catalyze monomethylation and symmetric dimethylation of arginine residues. PRMT7 as the only type III PRMT catalyzes monomethylation of arginine residues. All PRMTs can methylate multiple protein substrates and some PRMTs share the same substrates. For example, H4R3 can be methylated by PRMT1, PRMT5, PRMT6 and PRMT7,^6-9^ TP53 can be methylated by PRMT3 and PRMT5,^10, 11^ and H3R2 can be methylated by PRMT5, PRMT6 and PRMT7.^5, 12^ To further understand roles of each PRMT in cell biology, highly selective and cell-active inhibitors of each PRMT are needed. In addition to being invaluable chemical tools, selective PRMT inhibitors could also serve as leads for drug discovery programs. To date, highly selective inhibitors for PRMT3,^13^ CARM1,^14, 15^ PRMT5^16-20^ and PRMT7^21^ have been reported. However, it remains challenging to develop highly selective inhibitors for other PRMTs including PRMT6 due to high homology in the conserved catalytic core shared among type I PRMTs.^22^ PRMT6 is associated with proliferative phenotypes,^23^ and contributes to negative regulation of global DNA methylation in cancer cells.^24^ However, its function is still poorly understood. While several PRMT6 inhibitors have been reported to date, a highly selective and cell-active PRMT6 inhibitor that can be utilized as a PRMT6 chemical probe has not yet been reported. Previously, we reported the type I PRMT pan inhibitor MS023,^25^ PRMT6 covalent inhibitor MS117 with modest selectivity^26^ and PRMT4/6 dual inhibitor MS049.^25^ In addition, EPZ020411^27^ was reported as a PRMT6 inhibitor with limited selectivity. Here, we report the first highly selective and cell-active allosteric inhibitor of PRMT6, SGC6870, which is one of only few PRMT allosteric inhibitors including our previously reported PRMT3 allosteric inhibitor, SGC707,^13^ and a recently reported PRMT5 allosteric inhibitor^28^. We also developed SGC6870N, which is the enantiomer of SGC6870 but is inactive for PRMT6, as a negative control of SGC6870 for chemical biology studies.

## RESULTS and DISCUSSION

### Discoveries of PRMT6 SGC6870 and SGC6870N

Our quest for the discovery of a selective PRMT6 inhibitor began with the screening of a diverse library of 5,000 compounds, from which a submicromolar inhibitor (±)-**1** (IC_50_ = 755 ± 87 nM) was identified (Figure 1A). Optically pure (*R*)-**1** enantiomer and (*S*)-**1** enantiomer were further chirally separated. The (*R*)-**1** enantiomer was identified as the active isomer (IC_50_ = 388 ± 77 nM), while the (*S*)-**1** enantiomer was inactive (IC_50_ > 100 μM) (Figure 1A). To discover the first highly selective chemical probe of PRMT6, we conducted structure-activity relationship (SAR) studies and successively optimized the amide group, benzodiazepine ring substituent, and pendant ring moiety of compound (±)-**1**. Design, synthesis and testing of more than 60 derivatives of (±)-**1** led to the identification of compound (±)-**2**, which contains 3,5-dimethyl phenyl and thiophene rings, with the most potent inhibitory effect of PRMT6 (IC_50_ = 214 ± 37 nM). Optically pure (*R*)-enantiomer SGC6870 and (*S*)-enantiomer SGC6870N were then obtained by chiral separation. Similar to compound (±)-**1,** the (*R*)-enantiomer of (±)-**2**, SGC6870, showed potent PRMT6 inhibition (IC_50_ = 77 ± 6 nM) while the (*S*)-enantiomer, SGC6870N, was inactive (IC_50_ > 50 μM) (Figure 1A, Figure 1B).

**Figure 1.**
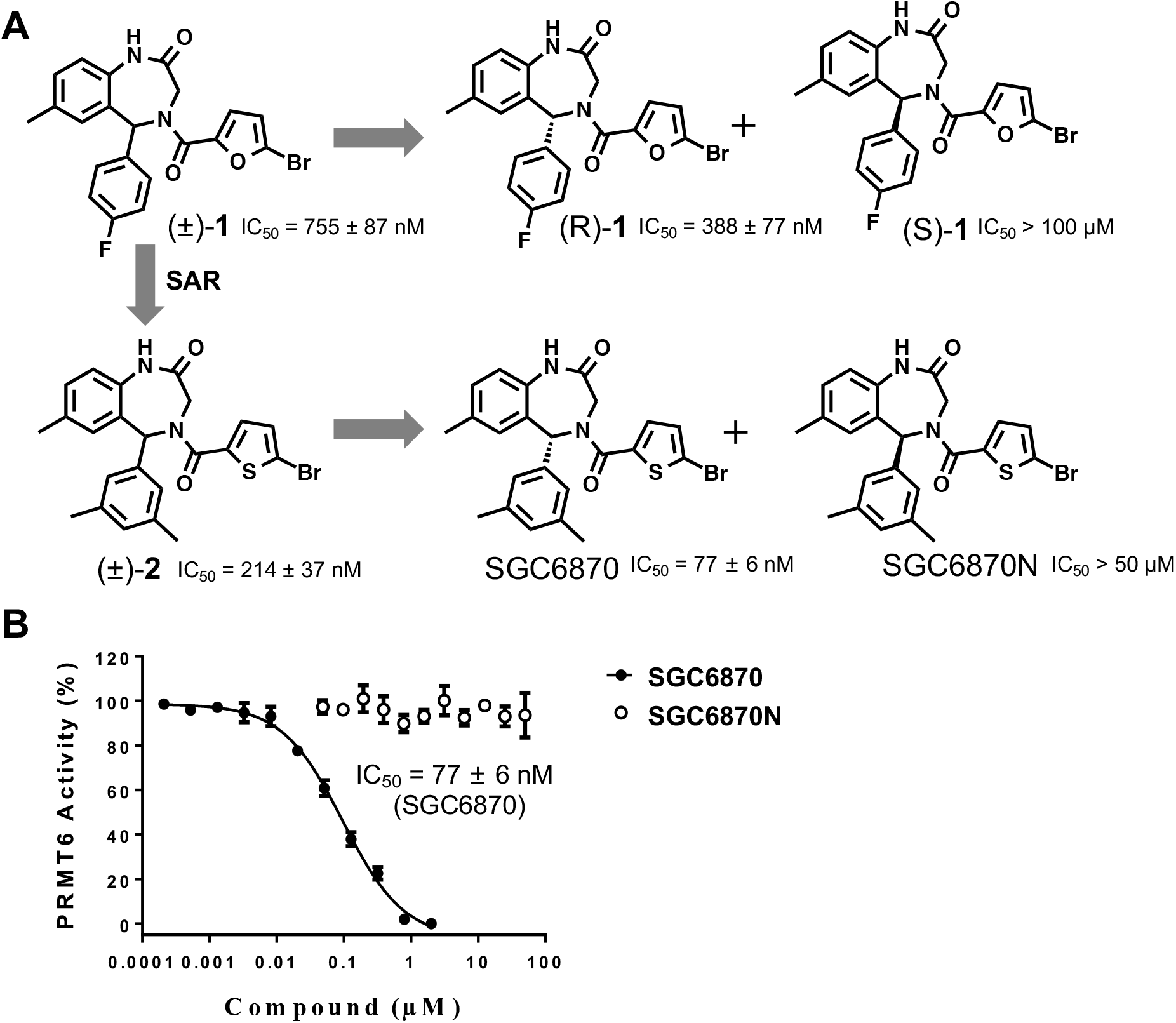
Discovery of the PRMT6 chemical probe SGC6870. (A) Structure-activity relationship studies and chiral separation led to the discovery of SGC6870 as a potent PRMT6 inhibitor and its enantiomer, SGC6870N, as a negative control. (B) IC_50_ determination of SGC6870 and SGC6870N, n = 3.

### Selectivity assessment

Next, we assessed selectivity of SGC6870 and SGC6870N against a total of 33 methyltransferases, including 8 PRMTs, 21 protein lysine methyltransferases (PKMTs), 3 DNA methyltransferases (DNMTs) and 1 RNA methyltransferase. SGC6870 at both 1 and 10 μM potently inhibited PRMT6, but did not significantly inhibit other 32 methyltransferases (Figure 2, Table S1). As expected, negative control SGC6870N did not significantly inhibit any of the 33 methyltransferases including PRMT6, at 1 or 10 μM (Figure 2, Table S1). Furthermore, SGC6870 was remarkably selective for PRMT6 over a broad range of non-epigenetic targets including kinases, G protein-coupled receptors (GPCRs), ion channels, and transporters at 1 μM (Table S2).

**Figure 2.**
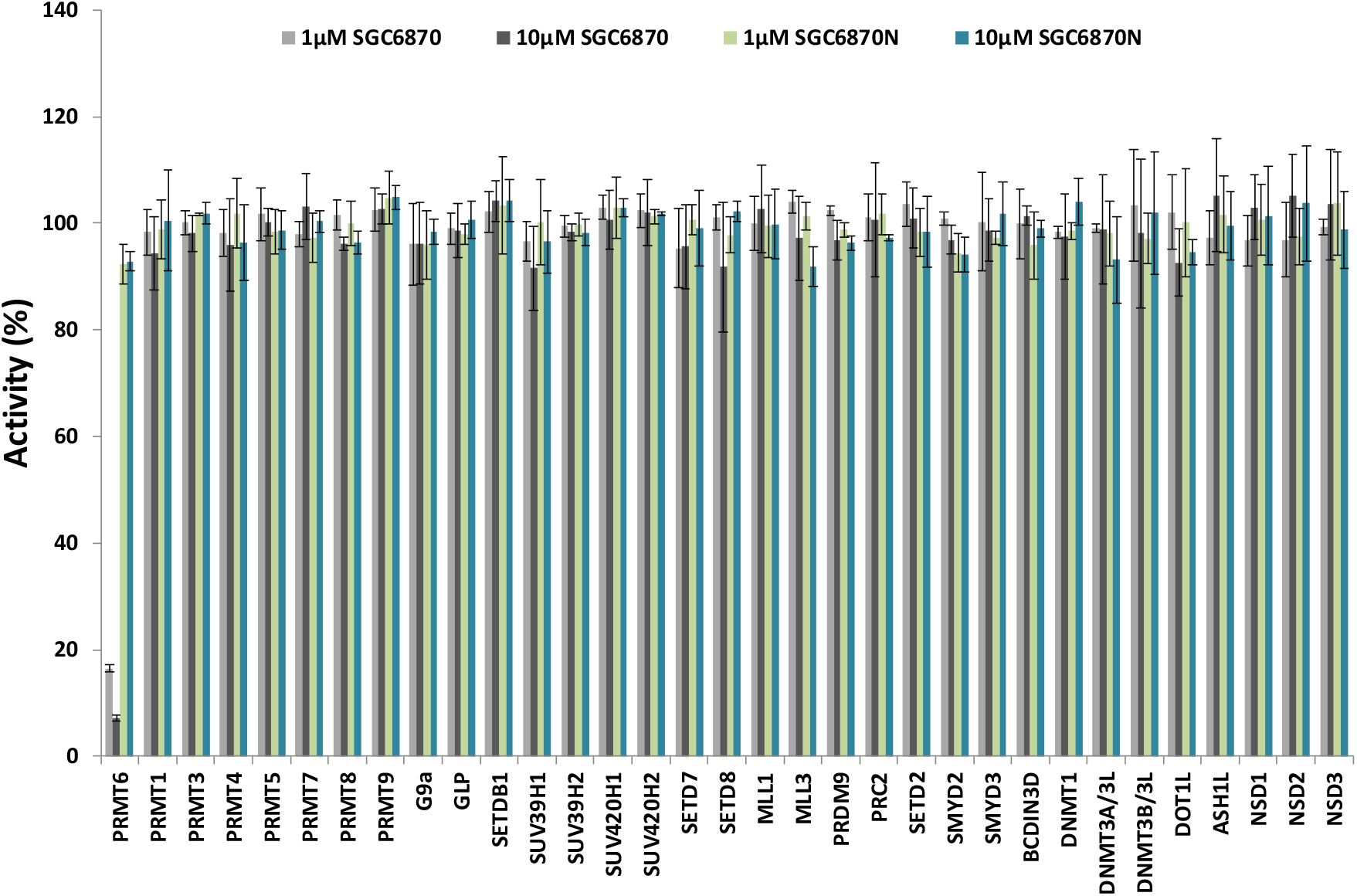
Selectivity assessment of SGC6870 at 1 μM (gray) and 10 μM (black) and SGC6870N at 1 μM (green) and 10 μM (blue) against 8 PRMTs, 21 protein lysine methyltransferases (PKMTs), 3 DNA methyltransferases (DNMTs) and 1 RNA methyltransferase, n = 3.

### Mechanism of action

Typically, when the addition of an inhibitor leads to quick equilibrium, the IC_50_ values can be determined very reproducibly regardless of small variations in preparation of reaction mixtures, and time of inhibitor incubation with protein. However, as we characterized SGC6870 inhibition, we observed significant changes in IC_50_ values we obtained from various experiments. We then investigated a possible time-dependent inhibition of PRMT6 by SGC6870 (Table S3). Inhibitory effect of SGC6870 was indeed time-dependent with more than 100-fold decrease in IC_50_ value after 2 hours of compound pre-incubation with PRMT6 (Table S3). This indicates that SGC6870 binds slowly and requires significant time to reach equilibrium before steady state turnover initiated by the addition of the substrate (Figure 3A, 3B and Table S3). To confirm that the time-dependent inhibition of PRMT6 by SGC6870 is not due to covalent binding of the compound, we tested samples of PRMT6 incubated with SGC6870 for 2 hours by mass spectrometry. No covalent modification of PRMT6 by SGC6870 was observed (Figure S1). Using 2 hour pre-incubation, we investigated the mechanism of action of SGC6870 by determining the IC_50_ values at various concentrations of the cofactor (Figure 3C) or substrate (Figure 3D). Our data supported a noncompetitive pattern of inhibition with respect to both the SAM cofactor and peptide substrate as no changes in IC_50_ values were observed as the cofactor or substrate concentrations were increased (Figure 3C, 3D). These mechanism of action results suggest that SGC6870 may be an allosteric inhibitor binding away from the catalytic active site.

**Figure 3.**
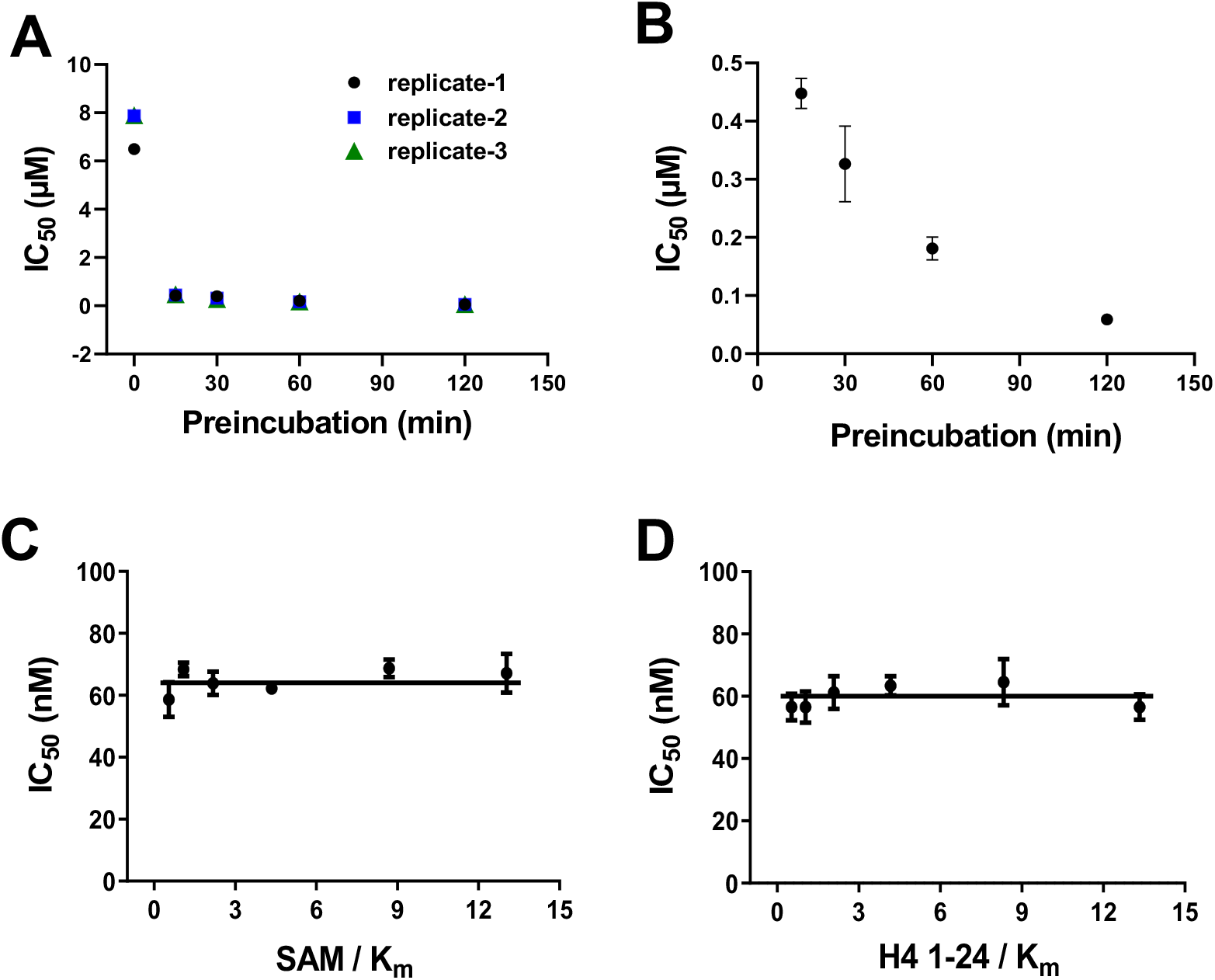
Mechanism of action. (A) IC_50_ values were determined at various compound-protein incubation times ranging from 15 to 120 minutes. The value obtained with no incubation time was used as a control (values are presented in Table S3). (B) Part of the plot in Figure C is magnified for better view of the decrease in IC_50_ value at longer incubation times. (C) IC_50_ values of SGC6870 at increasing concentrations of SAM (up to 30 μM) at fixed concentration of peptide substrate (3 μM of H4 1-24 peptide), and (D) IC_50_ values of SGC6870 at increasing concentration of peptide (up to 8 μM) at fixed concentration of SAM (12 μM).

### Cocrystal Structures

To further investigate the mechanism of inhibition, we solved the crystal structures of PRMT6 in complex with SGC6870 (PDB: 6W6D) (Figure 4A, Table S4) and (*R*)-**1** (PDB: 5WCF) (Figure S2A, Table S4), respectively. Structural alignment of PRMT6-SGC6870-SAM complex, and previously reported PRMT6-MS023-SAH^25^ and PRMT6-SAH^29^ complexes revealed SGC6870 binds in a unique pocket distinct from either the substrate or SAM binding pocket (Figure 4A, 4B) consistent with our kinetic data. A nine-amino acids loop consisting of residues Gly158 through Met166 flipped towards and downsized the substrate-binding pocket, creating a new pocket for accommodating SGC6870 (Figure 4B). Notably, this loop movement was highly localized and no other large-scale changes were observed in the complex. Several key interactions were observed between SGC6870 and residues in this new site (Figure 4A). The endo-amide on the diazepine ring formed a hydrogen bond with the backbone nitrogen of Gly158 and the exo-amide formed a hydrogen bond with the backbone nitrogen of Gly160. In addition, the thiophene group formed a T-shaped *π-π* stacking interaction with Tyr159 and another T-shaped *π-π* stacking interaction was also observed between the dimethylphenyl group and Trp156. Furthermore, the methylphenyl portion of the benzodiazepine moiety binds in a hydrophobic pocket. Residue Ala321 is one of the key components creating the newly formed binding pocket for the bromothiophene group of SGC6870 and bromofuran group of (*R*)-**1** (Figure S2B). We mutated it to Ile, Gln and Met bearing bulkier side chains, respectively. While these directed mutations did not significantly impact the substrate or SAM binding affinity to PRMT6 or PRMT6 catalytic activity itself (Table S5), they did impair PRMT6 inhibitory potency of (*R*)-**1** by 4 – 12-fold (Table S6). It should be noted that, structural alignment of PRMT6-SGC6870 and PMRT6-(*R*)-**1** complexes revealed two inhibitors possess the same binding mode (Figure S2B). Taken together, these results show both SGC6870 and (*R*)-**1** bind to the induced PRMT6 allosteric pocket. As previously reported, electron donating or withdrawing substituents of aromatic rings impact the T-shaped *π-π* stacking interactions.^30^ Compared with (*R*)-**1**, the enhanced inhibitory potency of SGC6870 could be partially due to higher electron density of thiophene and dimethylphenyl groups of SGC6870 than furan and fluorophenyl of (*R*)-**1**, as a result of stronger T-shaped *π-π* stacking interactions between SGC6870 and PRMT6 residues Tyr159 and Trp156. Collectively, both structural and kinetic data consistently revealed that SGC6870 is an allosteric PRMT6 inhibitor.

**Figure 4.**
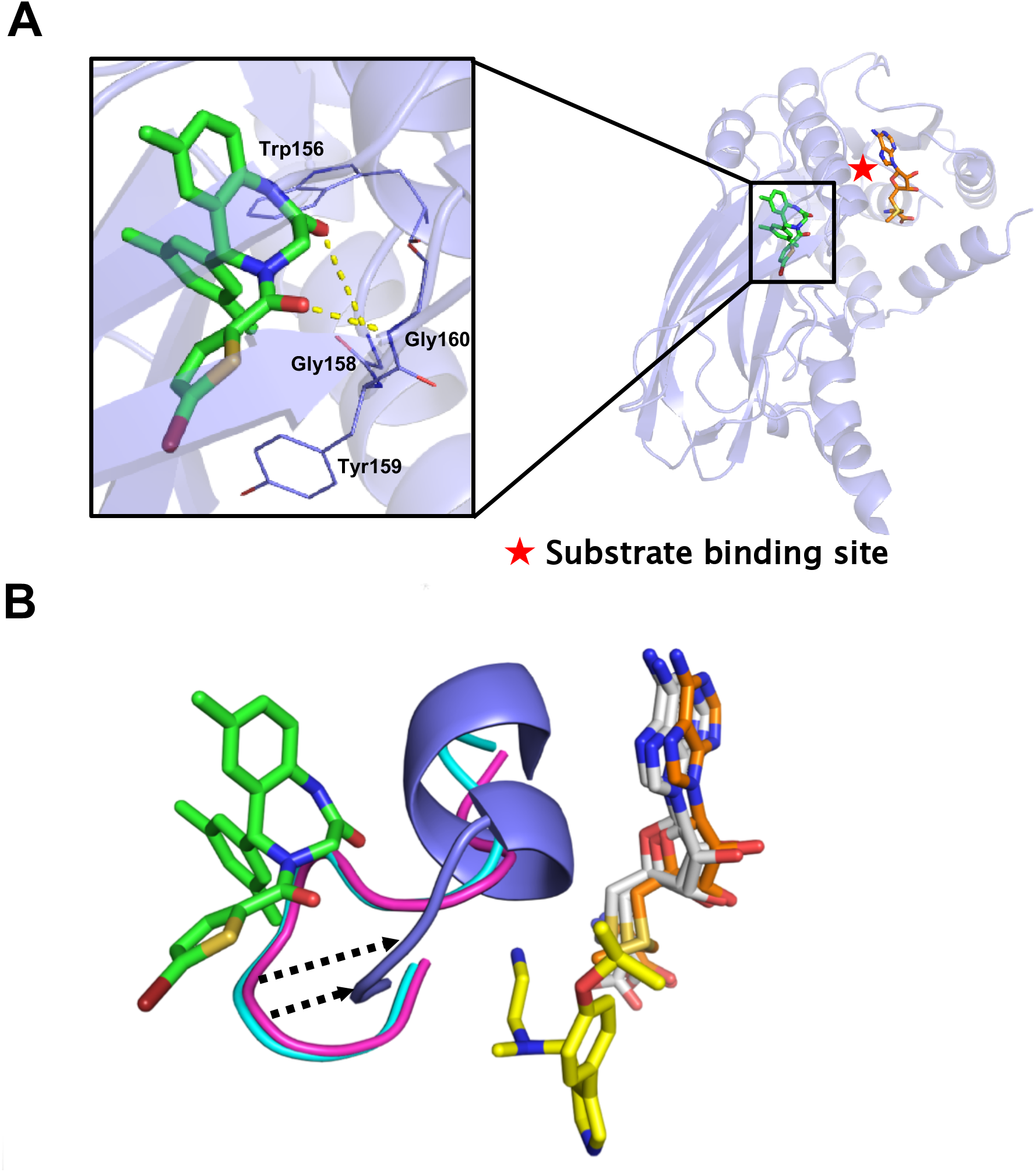
Cocrystal structure of PRMT6 in complex with SGC6870. (A) Cocrystal structure of PRMT6 (tinted blue) in complex with SGC6870 (green), and SAM (orange), (PDB: 6W6D). Dashed yellow lines indicate key hydrogen bonds. (B) Structural alignments of complexes PRMT6 (blue)-SGC6870 (green)-SAM (orange), PRMT6 (cyan)-MS023 (yellow)-SAH (gray) (PDB: 5E8R) and PRMT6 (magenta)-SAH (gray) (PDB: 4C05).

### Evaluation in Cellular Assays

Next, the cellular potency of SGC6870 against ectopically expressed PRMT6 in HEK293T cells was evaluated and a catalytically inactive PRMT6 mutant (V86K/D88A) was used as a positive control. HEK293T cells were treated with SGC6870 (Figure 5A) or SGC6870N (Figure 5B) at titrated concentrations for 20 h. SGC6870 potently and concentration-dependently reduced cellular levels of H3R2me2a (IC_50_ = 0.9 ± 0.1 μM) (Figure 5C) and H4R3me2a (IC_50_ = 0.6 ± 0.1 μM) (Figure 5D), both of which are known substrates of PRMT6.^5^ Furthermore, SGC6870 did not showed significant toxicity to HEK293T cells at up to 10 μM (Figure 5A). As expected, ectopical expression of the PRMT6 catalytically inactive V86K/D88A mutant led to near complete reduction of H3R2me2a and H4R3me2a markers (Figure 5A, 5B). Consistent with its poor potency against PRMT6 in the biochemical assay, SGC6870N did not significantly reduce cellular levels of H4R3me2a or H3R2me2a at up to 10 μM (Figure 5B). However, at 30 μM, significant toxicity to HEK293T cells was observed (Figure 5B). Finally, we further investigated the effect of SGC6870 and SGC6870N on cell growth in three different cell lines, HEK293T (embryonic kidney), PNT2 (prostate) and MCF-7 (breast cancer). Neither SGC6870 nor SGC6870N showed any significant toxicity to these three cell lines at concentrations up to 10 μM (Figure 5E, 5F). Therefore, SGC6870 and SGC6870N can be utilized in cellular studies at up to 10 μM concentration.

**Figure 5.**
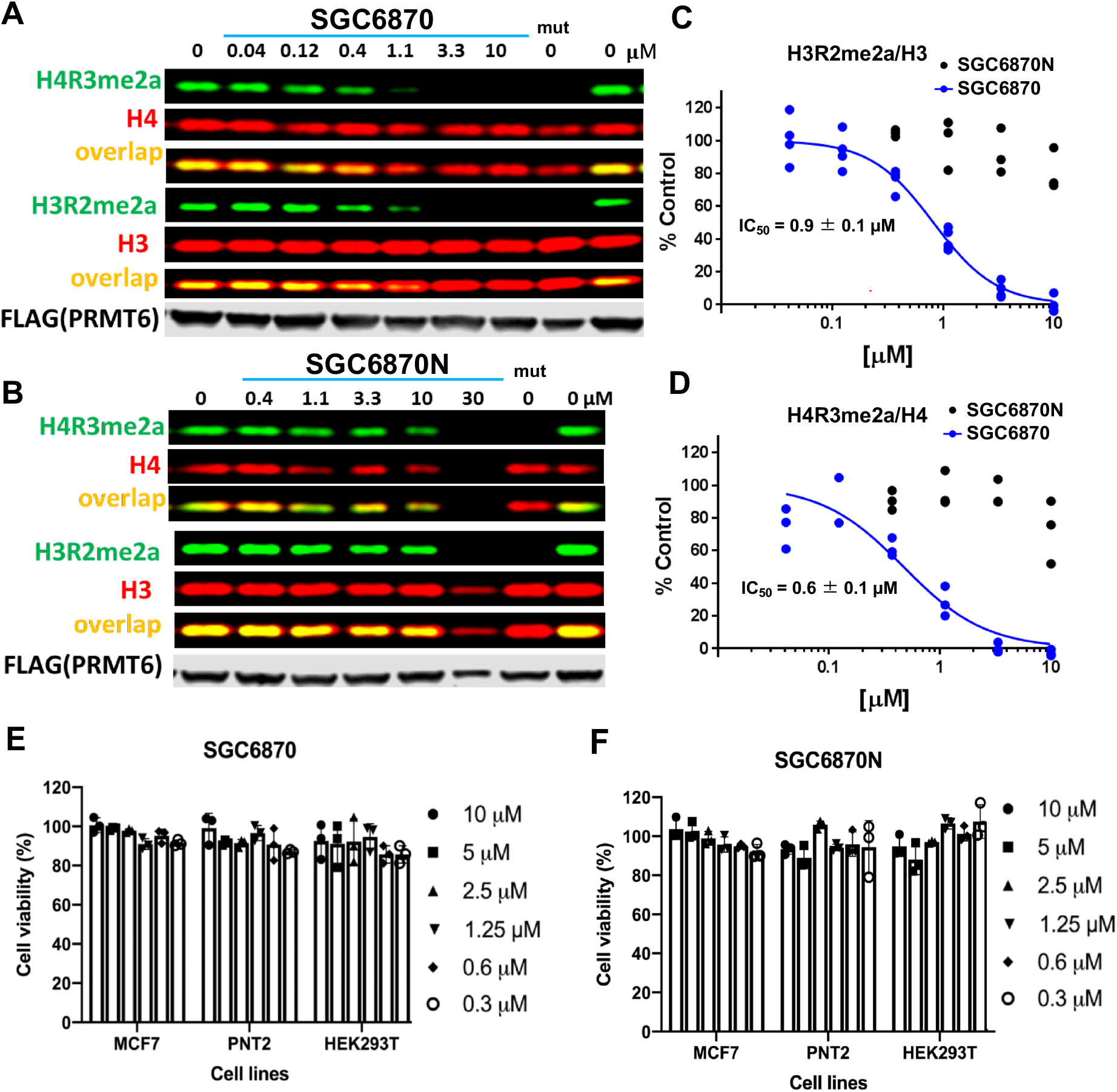
Inhibition of PRMT6 dependent H3R2 and H4R3 asymmetric di-methylation in cells. HEK293T cells were transfected with Flag-tagged PRMT6 and treated with indicated compounds for 20 h. The Flag-tagged PRMT6 catalytically inactive mutant V86K/D88A (mut) was used as a positive control. (A) Western blot representation of the effect of SGC6870 on PRMT6 activity. (B) Western blot representation of the effect of SGC6870N on PRMT6 activity. (C) IC_50_ determination of SGC6870 and SGC6870N at reducing H3R2me2a. The graph represents nonlinear fit of H3R2me2a fluorescence intensities normalized to intensities of H3, n = 4. (D) IC_50_ determination of SGC6870 and SGC6870N at reducing H4R3me2a. The graph represents nonlinear fit of H4R3me2a fluorescence intensities normalized to intensities of H4, n = 3. (E) Effect of SGC6870 on MCF-7, PNT2 and HEK293T cell viability, n = 3. (F) Effect of SGC6870N on MCF-7, PNT2 and HEK293T cell viability, n = 3.

## CONCLUSIONS

In conclusion, we discovered the first highly selective, potent and cell-active PRMT6 allosteric inhibitor, SGC6870. The crystal structure of the PRMT6-SGC6870 complex and kinetic studies revealed that SGC6870 binds to a unique, induced allosteric pocket, rendering this inhibitor highly selective for PRMT6 over 32 other methyltransferases, as well as a broad range of common drug targets. In addition, SGC6870 can significantly inhibit PRMT6 activity in cells with submicromolar potency, but did not display significant cellular toxicity at concentrations up to 10 μM. Furthermore, we demonstrated that the enantiomer of SGC6870, SGC6870N, showed very poor inhibitory effect on PRMT6 activity in both biochemical and cell-based assays and can be used as a negative control. Collectively, SGC6870 is a valuable chemical probe of PRMT6 and together with SGC6870N is an excellent tool compound to further investigate physiological and pathophysiological functions of PRMT6.

## ASSOCIATED CONTENT

### Supporting information

The Supporting Information is available free of charge at https://pubs.acs.org Selectivity data, mass spectrometry data, effect of preincubation on PRMT6 inhibitory potency of SGC6870, activity assessment on PRMT6-wild type (1-375) and its mutants, inhibitory potency of (*R*)-**1** against PRMT6-wild type (1-375) and its mutants, crystal structure of PRMT6 with (*R*)-**1**, detailed experimental procedures, crystallographic statistics and collection parameters, synthesis and spectral characterization of compounds (PDF)

## Author Contributions

## ACKNOWLEDGMENTS

This work was supported in part by the grants R01GM122749 (to J.J.) and P30CA196521 (to J.J.) from the U.S. National Institutes of Health and an endowed professorship from the Icahn School of Medicine at Mount Sinai (to J.J.). The SGC is a registered charity (number 1097737) that receives funds from AbbVie, Bayer Pharma AG, Boehringer Ingelheim, Canada Foundation for Innovation, Eshelman Institute for Innovation, Genome Canada through Ontario Genomics Institute [OGI-055], Innovative Medicines Initiative (EU/EFPIA) [ULTRA-DD grant no. 115766], Janssen, Merck KGaA, Darmstadt, Germany, MSD, Novartis Pharma AG, Ontario Ministry of Research, Innovation and Science (MRIS), Pfizer, São Paulo Research Foundation-FAPESP, Takeda, and Wellcome [106169/ZZ14/Z]. Eli Lilly and Company was a funder of SGC during the discovery and characterization of (*R*)-**1**. This work utilized the AVANCE NEO 600 MHz NMR Spectrometer System that was upgraded with funding from a National Institutes of Health SIG grant 1S10OD025132-01A1. We also thank Alice Shi Ming Li for contribution to biological chaterization of compounds.

## Table of Content Graphic

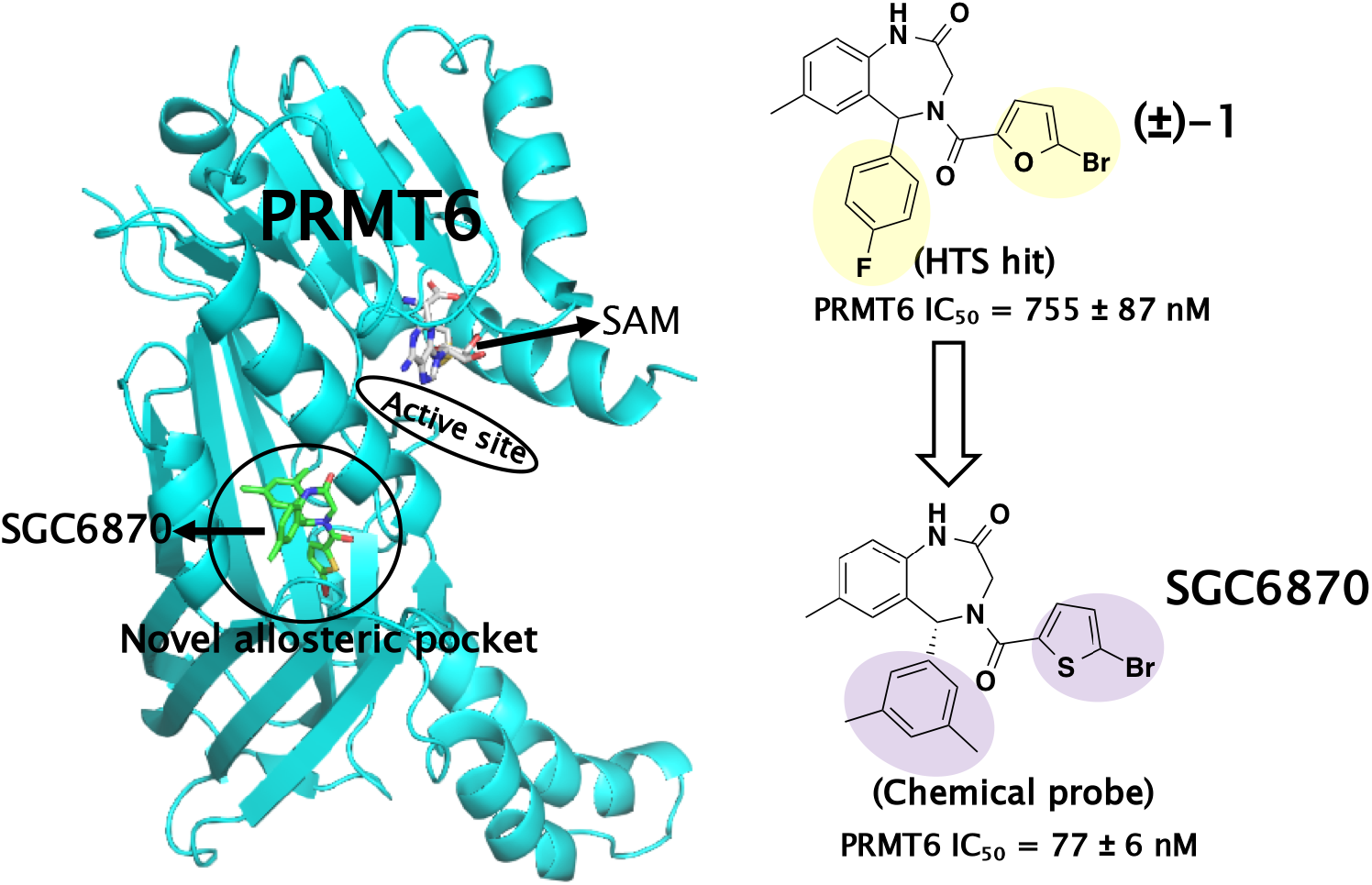

