## supporting information for "A First-in-class, Highly Selective and Cell-active Allosteric Inhibitor of Protein Arginine Methyltransferase 6 (PRMT6)"

**Table S1.** Selectivity of SGC6870 and SGC6870N against methyltransferases. 2

**Table S2.** Selectivity of SGC6870 against non-epigenetic targets. 3

**Table S3.** Effect of preincubation on PRMT6 inhibitory potency of SGC6870. 4

**Table S4.** Crystallography data and refinement statistics. 5

**Table S5.** Activity assessment on PRMT6-wild type (1-375) and its mutants. 6

**Table S6.** Inhibitory potency of (*R*)-**1** against PRMT6-wild type (1-375) and its mutants. 7

**Figure S1.** Mass spectrometry results of PRMT6 incubated with SGC6870. 8

**Figure S2.** Crystal structures of PRMT6 in complex with an allosteric inhibitor. 9

**Figure S3.** $^{1}$H NMR spectrum of compound 3. 10

**Figure S4.** $^{13}$C NMR spectrum of compound 3. 10

**Figure S5.** $^{1}$H NMR spectrum of SGC6870. 11

**Figure S6.** $^{13}$C NMR spectrum of SGC6870. 11

**Figure S7.** $^{1}$H NMR spectrum of SGC6870N. 12

**Figure S8.** $^{13}$C NMR spectrum of SGC6870N. 12

**Supplementary Experimental Section** 13

**Table S1.** Selectivity of SGC6870 and SGC6870N against methyltransferases

| Protein | Activity% | | | | | | | | | | | |
|---|---|---|---|---|---|---|---|---|---|---|---|---|
| | SGC6870 | | | | | | SGC6870N | | | | | |
| | 1μM | | | 10μM | | | 1μM | | | 10μM | | |
| | exp1 | exp2 | exp3 | exp1 | exp2 | exp3 | exp1 | exp2 | exp3 | exp1 | exp2 | exp3 |
| PRMT6 | 16 | 17 | 17 | 8 | 7 | 7 | 90 | 97 | 90 | 93 | 91 | 94 |
| PRMT1 | 103 | 98 | 94 | 102 | 91 | 90 | 105 | 97 | 94 | 92 | 110 | 100 |
| PRMT3 | 101 | 102 | 98 | 102 | 95 | 97 | 102 | 102 | 101 | 102 | 100 | 104 |
| PRMT4 | 100 | 101 | 93 | 90 | 106 | 93 | 94 | 106 | 105 | 93 | 105 | 92 |
| PRMT5 | 101 | 98 | 107 | 98 | 103 | 99 | 99 | 94 | 103 | 102 | 95 | 100 |
| PRMT7 | 99 | 99 | 95 | 106 | 107 | 96 | 102 | 93 | 98 | 101 | 102 | 98 |
| PRMT8 | 102 | 98 | 104 | 97 | 96 | 95 | 97 | 105 | 98 | 95 | 96 | 99 |
| PRMT9 | 107 | 100 | 100 | 102 | 106 | 100 | 107 | 108 | 99 | 107 | 103 | 105 |
| G9a | 91 | 92 | 105 | 105 | 91 | 93 | 102 | 96 | 89 | 101 | 98 | 96 |
| GLP | 101 | 100 | 96 | 104 | 94 | 99 | 100 | 98 | 96 | 99 | 99 | 105 |
| SETDB1 | 98 | 105 | 104 | 108 | 100 | 104 | 110 | 107 | 93 | 109 | 102 | 102 |
| SUV39H1 | 101 | 95 | 94 | 100 | 90 | 84 | 109 | 96 | 95 | 93 | 103 | 94 |
| SUV39H2 | 101 | 97 | 100 | 98 | 97 | 100 | 99 | 102 | 98 | 100 | 99 | 95 |
| SUV420H1 | 105 | 102 | 101 | 106 | 96 | 100 | 96 | 106 | 107 | 103 | 105 | 101 |
| SUV420H2 | 99 | 105 | 103 | 109 | 98 | 99 | 100 | 103 | 101 | 101 | 102 | 102 |
| SETD7 | 97 | 102 | 87 | 102 | 98 | 87 | 101 | 103 | 98 | 106 | 99 | 92 |
| SETD8 | 104 | 100 | 99 | 104 | 93 | 79 | 100 | 100 | 94 | 100 | 103 | 104 |
| MLL1 | 95 | 105 | 99 | 112 | 97 | 99 | 98 | 106 | 94 | 105 | 93 | 102 |
| MLL3 | 102 | 103 | 107 | 95 | 90 | 106 | 98 | 103 | 103 | 88 | 95 | 93 |
| PRDM9 | 103 | 101 | 103 | 94 | 96 | 101 | 100 | 97 | 99 | 98 | 96 | 95 |
| PRC2 | 96 | 102 | 105 | 109 | 104 | 89 | 105 | 102 | 98 | 97 | 97 | 98 |
| SETD2 | 99 | 107 | 105 | 107 | 96 | 99 | 103 | 94 | 97 | 91 | 104 | 100 |
| SMYD2 | 102 | 100 | 101 | 100 | 96 | 94 | 98 | 94 | 91 | 92 | 93 | 98 |
| SMYD3 | 104 | 107 | 90 | 105 | 95 | 95 | 97 | 96 | 99 | 101 | 108 | 96 |
| BCDIN3D | 101 | 106 | 93 | 101 | 103 | 100 | 89 | 100 | 99 | 98 | 99 | 101 |
| DNMT1 | 99 | 99 | 97 | 107 | 93 | 93 | 100 | 99 | 97 | 101 | 102 | 109 |
| DNMT3A/3L | 99 | 100 | 98 | 111 | 92 | 94 | 104 | 99 | 92 | 90 | 87 | 102 |
| DNMT3B/3L | 108 | 91 | 111 | 109 | 83 | 103 | 102 | 98 | 92 | 97 | 94 | 115 |
| DOT1L | 109 | 95 | 103 | 91 | 100 | 87 | 111 | 91 | 98 | 93 | 93 | 97 |
| ASH1L | 99 | 92 | 101 | 112 | 93 | 110 | 104 | 108 | 94 | 97 | 95 | 107 |
| NSD1 | 102 | 93 | 96 | 106 | 96 | 107 | 106 | 102 | 93 | 99 | 94 | 112 |
| NSD2 | 97 | 104 | 90 | 110 | 96 | 109 | 104 | 94 | 95 | 100 | 95 | 116 |
| NSD3 | 101 | 98 | 99 | 104 | 93 | 114 | 114 | 95 | 102 | 92 | 99 | 106 |

**Table S2.** Selectivity of SGC6870 against non-epigenetic targets.

| Target | Percentage of Inhibition at 1 µM (%) | Positive Control |
|---|---|---|
| A2A (h) (agonist radioligand) | 2 | NECA |
| Alpha 1A (h) (antagonist radioligand) | 4 | WB4101 |
| Alpha 2A (h) (antagonist radioligand) | -8 | yohimbine |
| Beta 1 (h) (agonist radioligand) | -13 | atenolol |
| Beta 2 (h) (antagonist radioligand) | -7 | ICI118551 |
| BZD (central) (agonist radioligand) | -9 | diazepam |
| CB1 (h) (agonist radioligand) | -20 | CP55940 |
| CB2 (h) (agonist radioligand) | -19 | WIN 55212-2 |
| CCK1 (CCKA) (h) (agonist radioligand) | 1 | CCK-8s |
| D1 (h) (antagonist radioligand) | 9 | SCH23390 |
| D2S (h) (agonist radioligand) | -1 | 7-OH-DPAT |
| ETA (h) (agonist radioligand) | -24 | endothelin-1 |
| NMDA (antagonist radioligand) | 1 | CGS19755 |
| H1 (h) (antagonist radioligand) | 0 | pyrilamine |
| H2 (h) (antagonist radioligand) | -18 | cimetidine |
| MAO-A (antagonist radioligand) | 12 | clorgyline |
| M1 (h) (antagonist radioligand) | -2 | pirenzepine |
| M2 (h) (antagonist radioligand) | -5 | methoctramine |
| M3 (h) (antagonist radioligand) | 12 | 4-DAMP |
| N neuronal alpha 4beta 2 (h) (agonist radioligand) | -18 | nicotine |
| Delta (DOP) (h) (agonist radioligand) | -20 | DPDPE |
| Kappa (h) (KOP) (agonist radioligand) | 20 | U50488 |
| Mu (MOP) (h) (agonist radioligand) | 4 | DAMGO |
| 5-HT1A (h) (agonist radioligand) | 5 | 8-OH-DPAT |
| 5-HT1B (h) (antagonist radioligand) | 6 | serotonine |
| 5-HT2A (h) (agonist radioligand) | 12 | (±)-DOI |
| 5-HT2B (h) (agonist radioligand) | 1 | (±)-DOI |
| 5-HT3 (h) (antagonist radioligand) | -5 | MDL72222 |
| GR (h) (agonist radioligand) | 15 | dexamethasone |
| AR (h) (agonist radioligand) | -7 | testosterone |
| V1a (h) (agonist radioligand) | -5 | [d(CH2)51,Tyr(Me)2]-AVP |
| Ca2+ channel (L, dihydropyridine site) (antagonist radioligand) | -8 | nitrendipine |
| Potassium Channel hERG (human)- [3H] Dofetilide | 0 | terfenadine |
| KV channel (antagonist radioligand) | -10 | alpha-dendrotoxin |
| Na+ channel (site 2) (antagonist radioligand) | -22 | veratridine |
| Norepinephrine transporter (h) (antagonist radioligand) | -4 | protriptyline |
| Dopamine transporter (h) (antagonist radioligand) | 6 | BTCP |
| 5-HT transporter (h) (antagonist radioligand) | -17 | imipramine |
| COX1(h) | 17 | diclofenac |
| COX2(h) | 8 | NS398 |
| PDE3A (h) | -11 | milrinone |
| PDE4D2 (h) | -12 | Ro20-1724 |
| Lck kinase (h) | 1 | staurosporine |
| Acetylcholinesterase (h) | 7 | galanthamine |

Results are from two duplicate experiments.

**Table S3**. Effect of preincubation on PRMT6 inhibitory potency of SGC6870.

| Preincubation (min) | PRMT6 + SGC6870 | |
|---|---|---|
| | $IC_{50}$ (μM) | Hill Slope |
| 0 | 7.4 ± 0.8 | 0.7 |
| 15 | 0.44 ± 0.03 | 1.2 |
| 30 | 0.33 ± 0.07 | 1.1 |
| 60 | 0.18 ± 0.02 | 1.5 |
| 120 | 0.06 ± 0.002 | 1.7 |

**Table S4**. Crystallography data and refinement statistics.

| | **PRMT6 + (*R*)-1** | **PRMT6 + SGC6870** |
|---|---|---|
| **PDB Code** | 5WCF | 6W6D |
| **Data collection** | | |
| Space group | $I4_1$ | $I4_1$ |
| Cell dimensions | | |
| *a*, *b*, *c* (Å) | 94.7, 94.7, 108.7 | 94.6,94.6,108.1 |
| $\alpha$, $\beta$, $\gamma$ (°) | 90.00,90.00,90.00 | 90.00,90.00,90.00 |
| Resolution (Å) (highest resolution shell) | 50.00-1.98(2.01-1.98) | 50.00-1.91(1.94-1.91) |
| Measured reflections | 253023 | 230004 |
| Unique reflections | 33067 | 36823 |
| $R_{merge}$ | 6.0(90.7) | 7.6(97.6) |
| $I/\sigma I$ | 47.0(2.3) | 35.4(1.7) |
| Completeness (%) | 100.0(100.0) | 99.8(99.9) |
| Redundancy | 7.7(7.7) | 6.2(6.1) |
| **Refinement** | | |
| Resolution (Å) | 50.00-1.98 | 47.35-1.91 |
| No. reflections (test set) | 32078(966) | 35321(1487) |
| $R_{work}$/ $R_{free}$ (%) | 20.0/20.8 | 18.1/20.9 |
| No. atoms | | |
| Protein | 2604 | 2563 |
| Co-factor | 26 | 26 |
| Compound | 28 | 29 |
| Water | 102 | 132 |
| B-factors (Å$^2$) | | |
| Protein | 46.4 | 40.9 |
| Co-factor | 49.8 | 39.4 |
| Compound | 40.8 | 32.0 |
| Water | 48.1 | 43.9 |
| RMSD | | |
| Bond lengths (Å) | 0.009 | 0.008 |
| Bond angles (°) | 1.372 | 1.446 |
| Ramachandran plot % residues | | |
| Favored | 97.7 | 96.7 |
| Additional allowed | 2.3 | 3.3 |
| Generously allowed | 0.0 | 0.0 |
| Disallowed | 0.0 | 0.0 |

**Table S5.** Activity assessment on PRMT6-wild type (1-375) and its mutants (17, 19, 20).

| Parameters | Wild-Type | PRMT6-17 | PRMT6-19 | PRMT6-20 |
|---|---|---|---|---|
| Mutation | - | A321I | A321Q | A321M |
| $K_m$ B-$H_4$(1-24), μM | 0.17 | 0.4 | 0.18 | 0.17 |
| $k_{cat}$, $h^{-1}$ | 42 | 85 | 20 | 29 |
| $K_m$ SAM, μM | 2 | 4 | 3 | 4 |

**Table S6.** Inhibitory potency of (*R*)-**1** against PRMT6-wild type (1-375) and its mutants (17, 19, 20).

| Protein | $IC_{50}$ (μM) | Hill Slope |
|---|---|---|
| PRMT6-Wt | 4 | 1.3 |
| PRMT6-17 | 46 | 1 |
| PRMT6-19 | 17 | 1.5 |
| PRMT6-20 | 41 | 0.8 |

The $IC_{50}$ values in Table S6 were generated using 20 min pre-incubation.

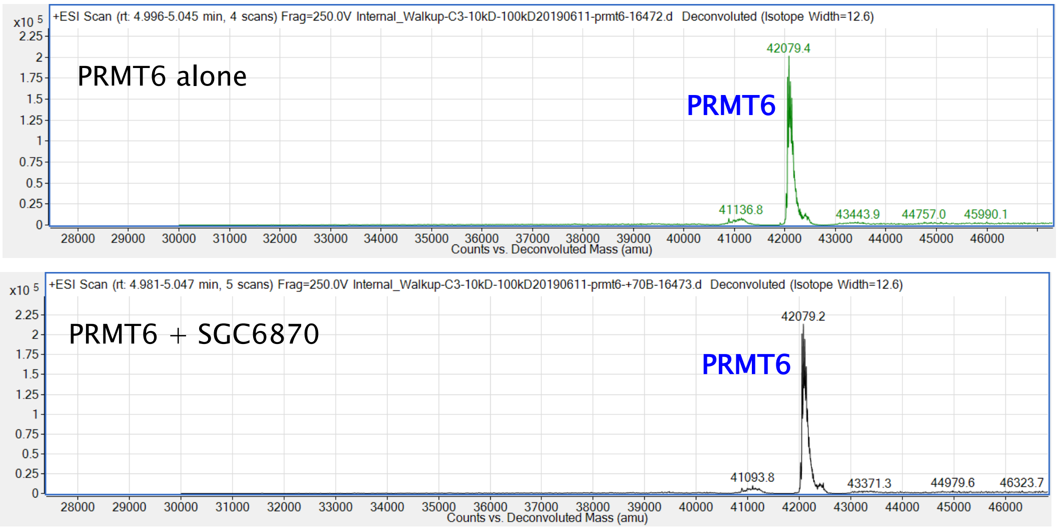


**Figure S1.** Mass spectrometry results of PRMT6 incubated with SGC6870 indicate that SGC6870 did not covalently modify PRMT6.

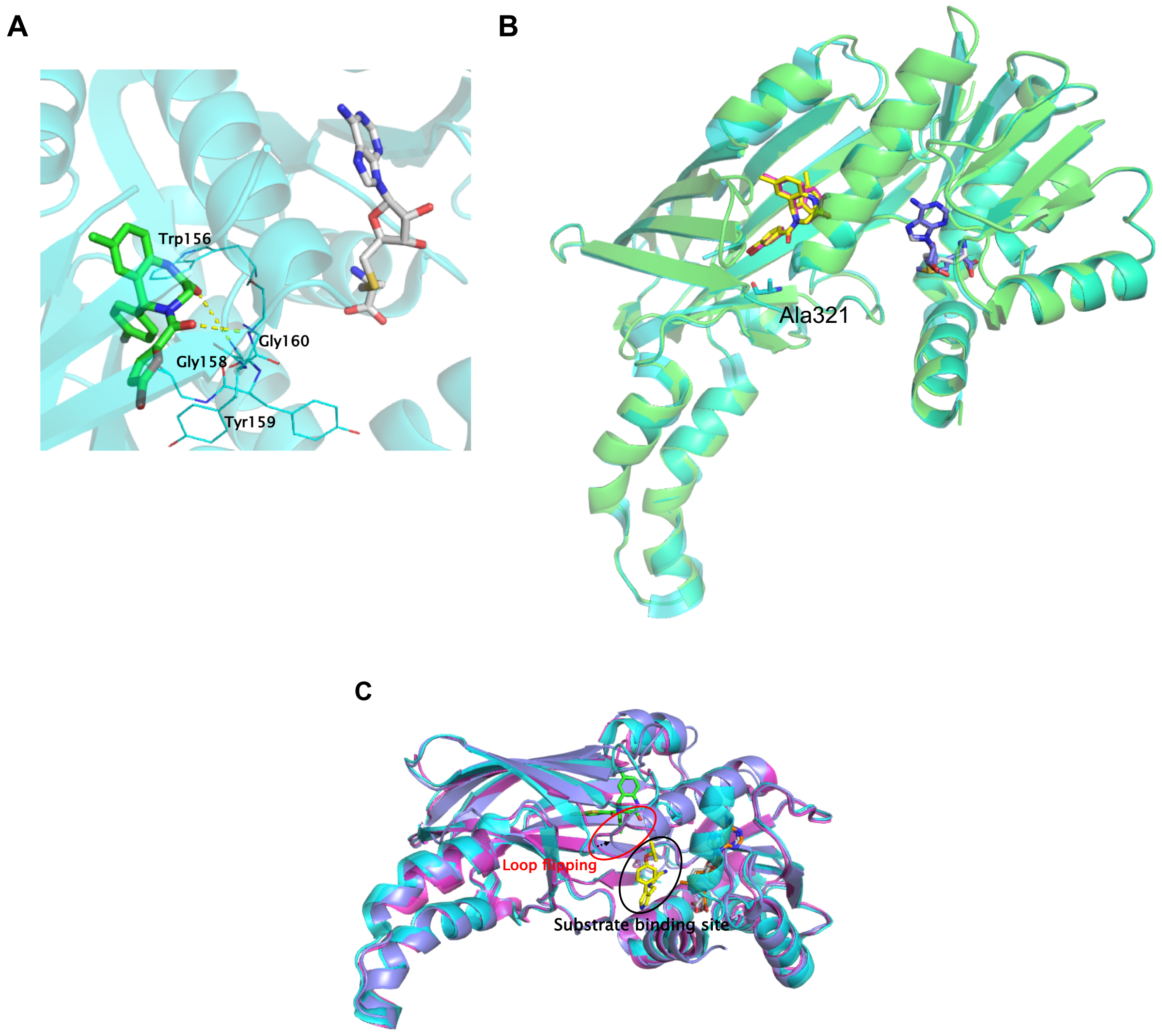


**Figure S2.** Crystal structures of PRMT6 in complex with an allosteric inhibitor. (A) Crystal structure of PRMT6 (cyan) in complex with (*R*)-**1** (green), SAH (gray), (PDB: 5WCF). Key hydrogen bond interactions are highlighted in yellow dotted lines. (B) Structural alignment of PRMT6 (cyan)-(*R*)-**1** (magenta)-SAH (gray) and PRMT6 (green)-SGC6870 (yellow)-SAM (blue). (C) Structural alignments of complexes PRMT6 (tint)-SGC6870 (green)-SAM (orange), PRMT6 (cyan)-MS023 (yellow)-SAH (gray) (PDB: 5E8R) and PRMT6 (magenta)-SAH (gray) (PDB: 4C05).

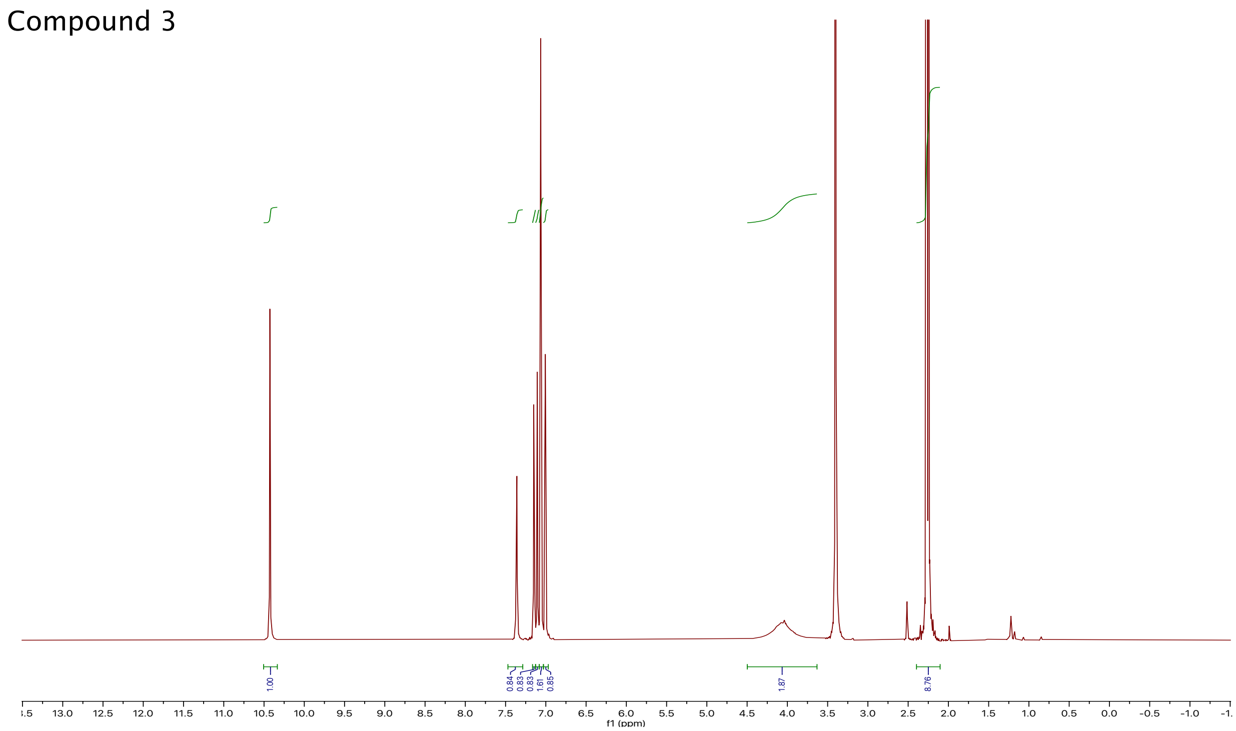


**Figure S3.** $^{1}H$ NMR spectrum of compound 3.

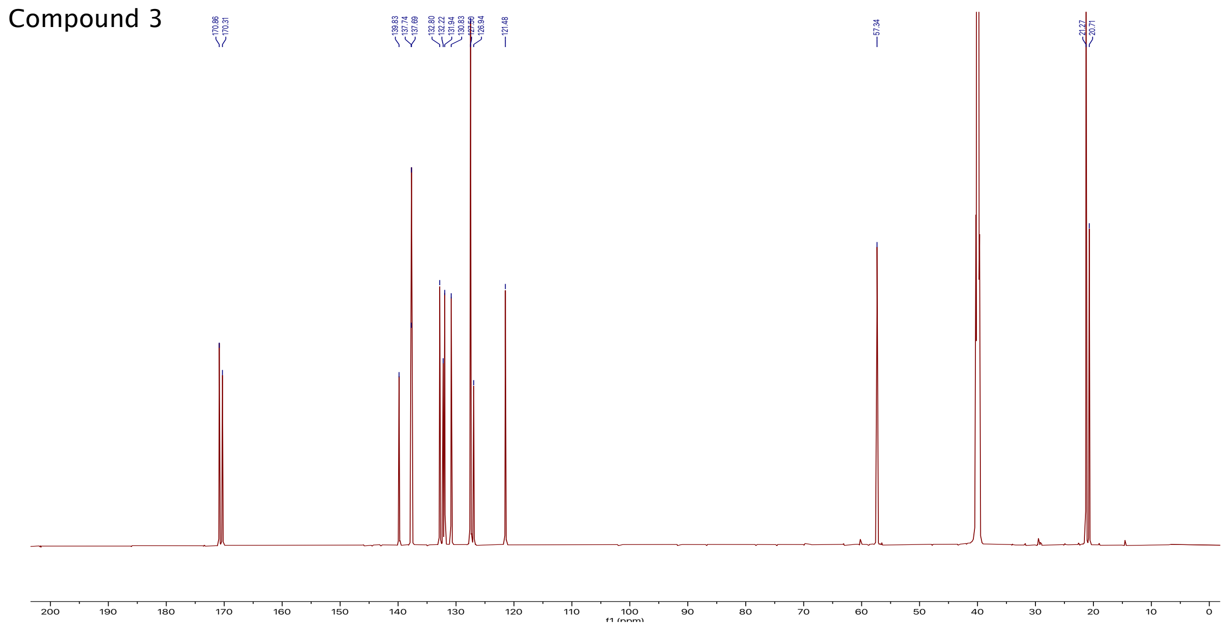


**Figure S4.** $^{13}C$ NMR spectrum of compound 3.

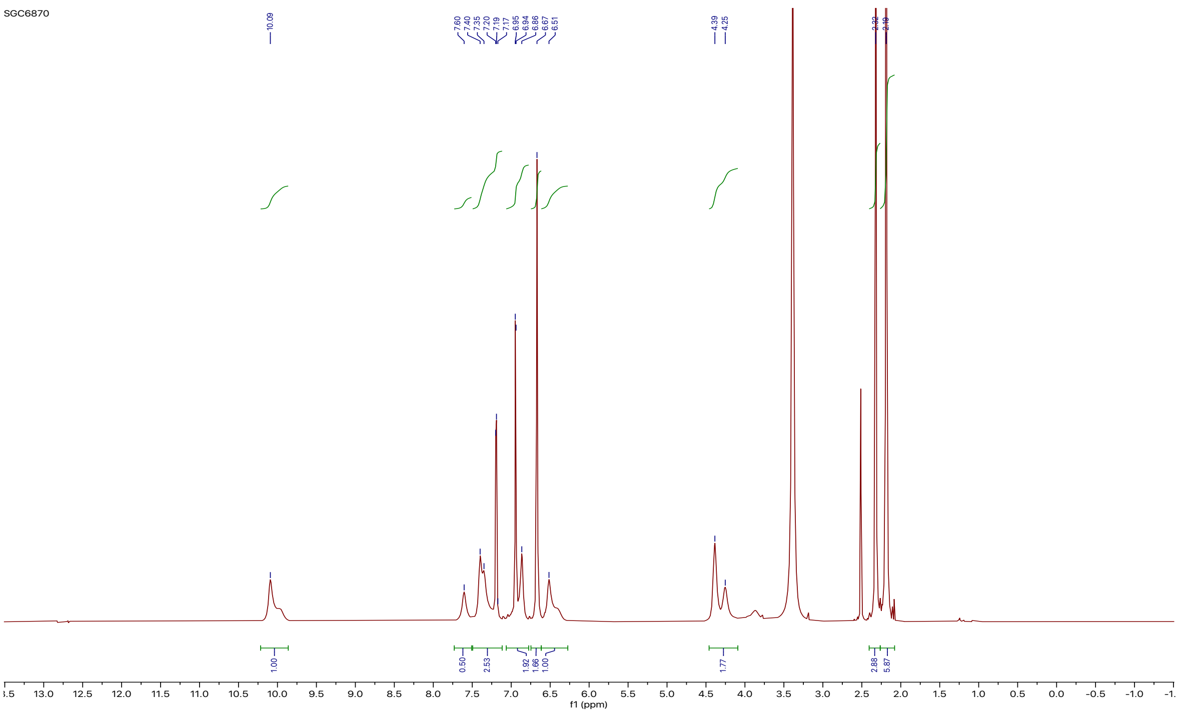

**Figure S5.** $^{1}H$ NMR spectrum of compound SGC6870.

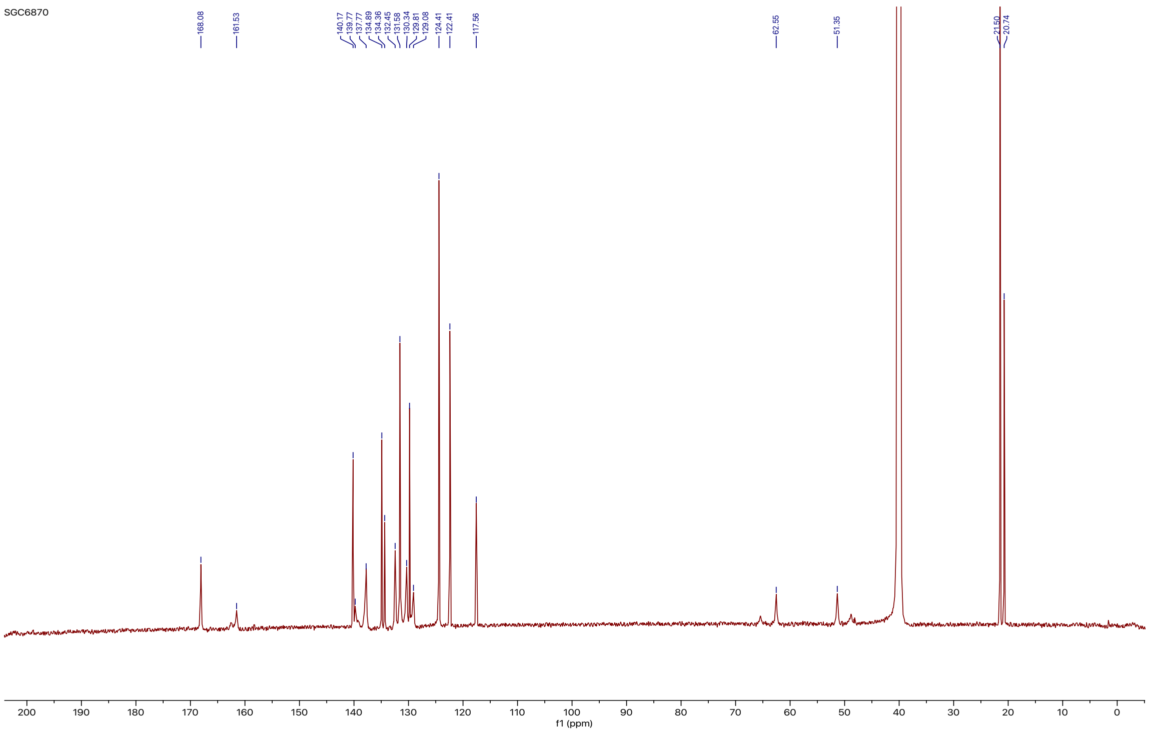

**Figure S6.** $^{13}C$ NMR spectrum of compound SGC6870.

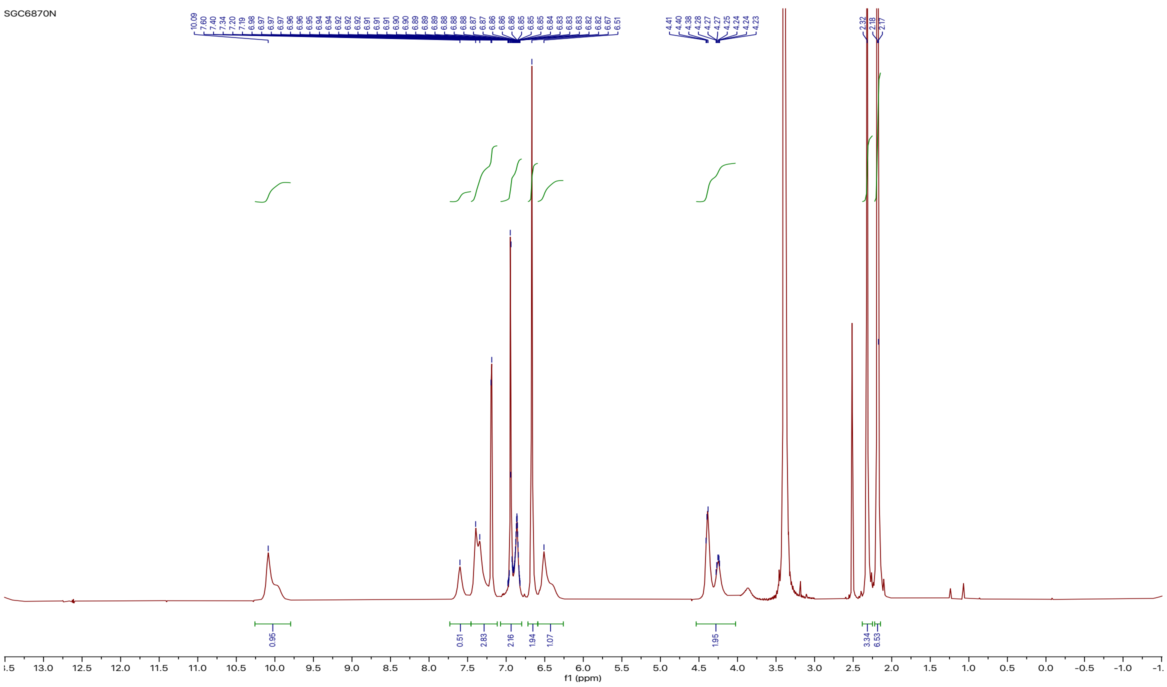

**Figure S7.** $^{1}$H NMR spectrum of compound SGC6870N.

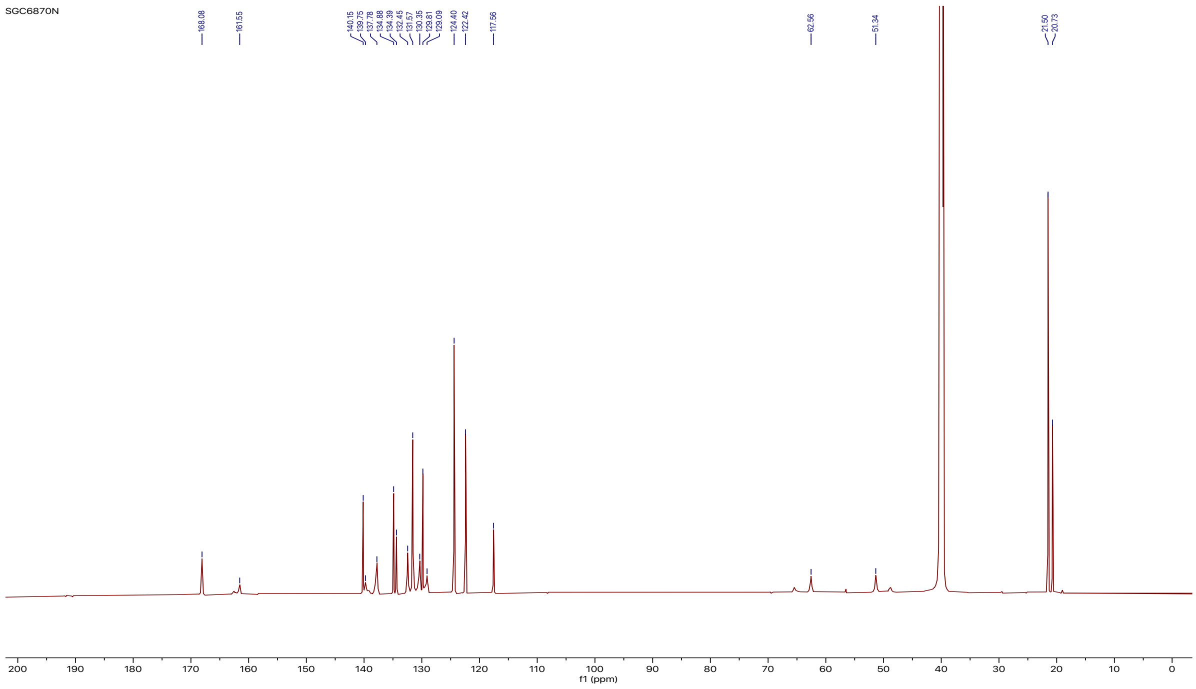

**Figure S8.** $^{13}$C NMR spectrum of compound SGC6870N.

**Supplementary Experimental Section**

**1. Synthetic procedures and compound characterization**

**Chemistry General Procedure.** All commercially available chemical reagents were directedly used without further purification. A Teledyne ISCO CombiFlash Rf+ instrument equipped with a variable wavelength UV detector and a fraction collector was used to conduct flash column chromatography. Redi*Sep*®*R*f HP C18 and flash silica columns were used for purification. High-resolution mass spectra (HRMS) data were acquired in positive ion mode using an Agilent G1969A API-TOF with an electrospray ionization (ESI) source. Nuclear Magnetic Resonance (NMR) spectra were acquired on Bruker Avance-III 800 MHz spectrometer (800 MHz $^1$H NMR, 201 MHz $^{13}$C NMR). Chemical shifts are reported in ppm (δ). The enantiomers were separated by prep SFC using an AS-H column (2 x 25 cm) and mobile phase of 25% MeOH/$CO_2$ with 0.1% DEA at 70 mL.min$^{-1}$ (100 bar).

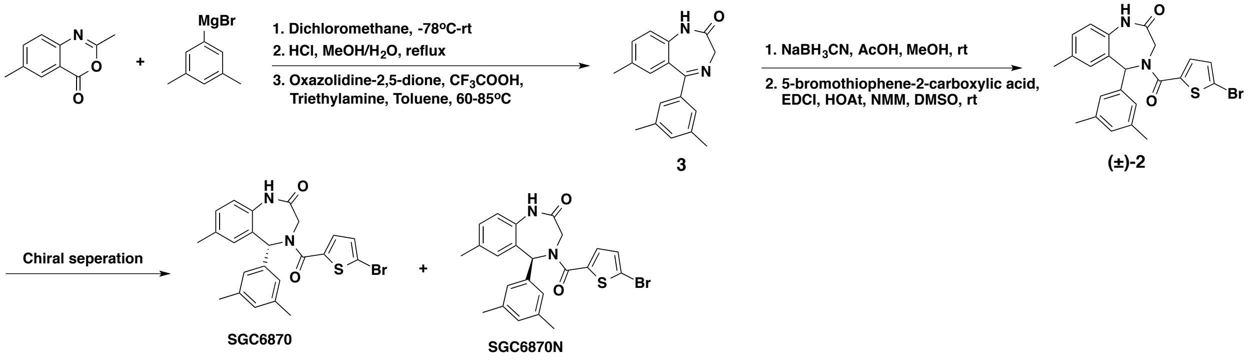


**5-(3,5-dimethylphenyl)-7-methyl-1,3-dihydro-2*H*-benzo[1,4]diazepin-2-one (3)** To the solution of 2,6-dimethyl-4*H*-benzo[*d*][1,3]oxazin-4-one (3.5 g , 20 mmol) in 60 mL of dichloromethane at -78ºC, was added dropwise 3,5-dimethylphenylmagnesium bromide (0.5 M in THF, 48 ml, 24 mmol) . The resulting mixture was slowly warmed to room temperature and stirred overnight. 30 mL of saturated aqueous ammonium chloride was carefully added and the mixture was extracted with dichloromethane (60 mL x 3). The organic phase was dried over anhydrous sodium sulfate and concentrated under reduced pressure. The resulting residue was dissolved in 60 mL of methanol and 10 mL of hydrochloric acid (36.5-38.0%). The solution was heated to reflux for 6h. The cooled solution was then carefully basified to pH = 8 by saturated aqueous sodium bicarbonate and concentrated under reduced pressure. The resulting mixture was extracted with ethyl acetate (50 mL x 3). The organic phase was dried over anhydrous sodium sulfate and concentrated under reduced pressure. The resulting residue was dissolved in 80 mL of toluene. To this solution, was added trifluoroacetic acid (1.7 mL, 22 mmol) and added portionwise oxazolidine-2,5-dione (2.6 g, 26 mmol). The reaction was heated to 60 ºC for 30 min. Then triethylamine (3.1 mL, 22 mmol) was added and the mixture was heated to 80 ºC for another 30 min. To the cooled mixture was carefully added 20 mL of saturated aqueous sodium bicarbonate and the mixture was extracted with ethyl acetate (50 mL x 3). The organic phase was dried over anhydrous sodium sulfate and concentrated under reduced pressure. The resulting residue was purified by flash chromatography on silica gel column with eluent (EtOAc/hexane: 0-50%) to give white solid (2.7 g, yield 49%). $^{1}$H NMR (800 MHz, DMSO-$d_6$) δ 10.43 (s, 1H), 7.36 (d, $J$ = 8.4 Hz, 1H), 7.15 (d, $J$ = 8.3 Hz, 1H), 7.11 (s, 1H), 7.06 (s, 2H), 7.01 (s, 1H), 4.46 – 3.66 (m, 2H), 2.27 (s, 6H), 2.25 (s, 3H). $^{13}$C NMR (201 MHz, DMSO-$d_6$) δ 170.86, 170.31, 139.83, 137.74, 137.69, 132.80, 132.22, 131.94, 130.83, 127.50, 126.94, 121.48, 57.34, 21.27, 20.71. MS (ESI) *m/z* 279.2 $[M+H]^+$. HRMS (TOF) *m/z* $[M + H]^+$ calcd for $C_{18}H_{19}N_2O^+$ 279.1492, found 279.1509.

**4-(5-bromothiophene-2-carbonyl)-5-(3,5-dimethylphenyl)-7-methyl-1,3,4,5-tetrahydro-2*H*-benzo[*e*][1,4]diazepin-2-one ((±)-2).** To the solution of 5-(3,5-dimethylphenyl)-7-methyl-1,3-dihydro-2*H*-benzo[*e*][1,4]diazepin-2-one (2.7 g, 9.7 mmol) in 40 mL of methanol, was added portionwise sodium cyanoborohydride (1.8 g, 28 mmol) and acetic acid (5.5 mL, 97 mmol). The mixture was stirred at room temperature overnight and the volatile was removed under reduced pressure. The residue was carefully treated by 20 mL of saturated aqueous sodium bicarbonate and extracted with ethyl acetate (50 mL x 3). The organic phase was dried over anhydrous sodium sulfate and concentrated under reduced pressure. To the solution of residue above in 50 mL of dimethyl sulfoxide, was added 5-bromothiophene-2-carboxylic acid (2 g, 10 mmol), EDCI (2.4 g, 13 mmol), HOAt (1.7 g, 13 mmol) and *N*-methylmorpholine (2.8 mL, 26 mmol). The resulting solution was stirred at room temperature for 8 h and extracted with ethyl acetate (150 mL) and water (100 mL). The organic phase was washed by another 100 mL of water for twice and concentrated. The residue was purified by flash chromatography on silica gel column with eluent (EtOAc/hexane: 0-50%) to give pale yellow solid (2.7 g, yield 59%).

The enantiomers were separated by prep SFC using an AS-H column (2 x 25 cm) and mobile phase of 25% MeOH/$CO_2$ with 0.1% DEA at 70 mL.min$^{-1}$ (100 bar). 15 mg aliquots in EtOH:DCM (1:1; 1 mL) were injected and absorbance monitored at 220 nm. Analytical SFC was used to determine the purity of combined separated fractions using an AS-H column (0.46 x 25 cm) and mobile phase of 40% MeOH/$CO_2$ with 0.1% DEA at 3 mL.min$^{-1}$ (120 bar). Peak 1 eluting at 2.44 min using the analytical conditions corresponded to SGC6870N, whereas Peak 2 eluting at 3.34 min corresponded to SGC6870. 1.0 g of each enantiomer was obtained from chiral separation of 2.2 g of racemic material.

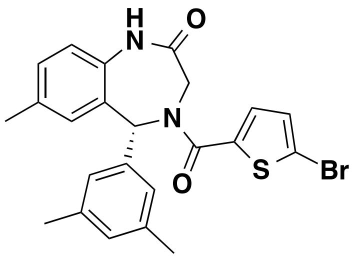


**SGC6870**

**SGC6870:** $^1$H NMR (800 MHz, DMSO-$d_6$) δ 10.19 – 9.82 (m, 1H), 7.73 – 7.49 (m, 1H), 7.41 – 7.33 (m, 2H), 7.19 (d, $J$ = 8.1 Hz, 1H), 7.00 – 6.81 (m, 2H), 6.67 (s, 2H), 6.59 – 6.31 (m, 1H), 4.48 – 3.82 (m, 2H), 2.32 (s, 3H), 2.19 (s, 6H). $^{13}$C NMR (201 MHz, DMSO-$d_6$) δ 168.08, 161.53, 140.17, 139.77, 137.77, 134.89, 134.36, 132.45, 131.58, 130.34, 129.81, 129.08, 124.41, 122.41, 117.56, 62.55, 51.35, 21.50, 20.74. MS (ESI) *m/z* 469.1 $[M+H]^+$. HRMS *m/z* $[M + H]^+$ calcd for $C_{23}H_{22}BrN_2O_2S^+$ 469.0580, found 469.0574. $[\alpha]_D^{20}$ +275.2° (*c* 4.55, MeOH).

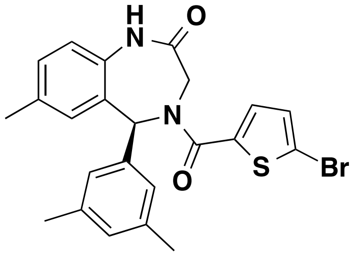


SGC6870N

**SGC6870N:** $^1$H NMR (800 MHz, DMSO-$d_6$) δ 10.22 – 9.85 (m, 1H), 7.60 (s, 1H), 7.49 – 7.10 (m, 3H), 7.01 – 6.80 (m, 2H), 6.67 (s, 2H), 6.60 – 6.27 (m, 1H), 4.55 – 3.75 (m, 2H), 2.32 (s, 3H), 2.18 (s, 6H). $^{13}$C NMR (201 MHz, DMSO-$d_6$) δ 168.08, 161.54, 140.15, 139.75, 137.78, 134.88, 134.39, 132.45, 131.57, 130.35, 129.81, 129.09, 124.40, 122.42, 117.56, 62.56, 51.34, 21.50, 20.73. MS (ESI) *m/z* 469.1 $[M+H]^+$. HRMS *m/z* $[M + H]^+$ calcd for $C_{23}H_{22}BrN_2O_2S^+$ 469.0580, found 469.0590. $[\alpha]_D^{20}$ -268.2° (*c* 4.77, MeOH).

Due to the amide rotamers of SGC6870 and SGC6870N, which can interconvert at ambient temperature, broadening of NMR peaks was observed.

### 2. Crystallization, Structure Determination

**Crystallization.** Human PRMT6 protein was expressed and purified according to the previously published protocol.[1] PRMT6 at 5.6 mg/mL was mixed with 5-fold molar excess of S-adenosyl-L-homocysteine (SAH) or S-Adenosyl methionine (SAM) and 3-fold molar excess of (*R*)-**1** or SGC6870 (dissolved from a previously prepared 100 mM DMSO stock solution). Diffraction quality crystals were obtained by setting a 96-well vapour-diffusion sitting drops at room temperature, in a precipitant solution containing 0.1 M sodium malonate pH 7.0, 12% (w/v) PEG 3350. The PRMT6- SGC6870/(*R*)-**1** crystal was cryo-protected by immersing it into a precipitant solution supplemented with 10% glycerol and then into paratone, and cryo-cooled in liquid nitrogen.

**Structure Determination.** X-ray diffraction data for PRMT6 + SGC6870/(*R*)-**1** were collected at 100K at beam line 24IDE of Advanced Photon Source (APS), Argonne National Laboratory and CMCF 08ID-1 of Canadian Light Source respectively. Both data sets were processed using the HKL-3000 suite.[2] The structures of PRMT6 + (*R*)-1 were directly refined against PDB entry 4HC4 as template. The structures of PRMT6 + SGC6870 were directly refined against PDB entry 5WCF as template. REFMAC was used for structure refinement.[3] GRADE was used to generate all restrains for compound refinement.[4] Graphics program COOT was used for model building and visualization.[5] Molprobity was used for structure validation.[6]

#### 3. PRMT6 Mutation

Four PRMT6 mutants were generated by mutating the alanine residue at position 321 of the wild type PRMT6 (PRMT6-WT) (residues 1-375) to isoleucine, arginine, glutamine and methionine of mutants 17, 19, and 20, respectively.

#### 4. Kinetic Characterization of PRMT6-WT and -mutants

Using the optimized assay conditions (25 nM of PRMT6-wt or-mutants, 20 mM Tris-HCl pH 7.5, 0.01% Triton X-100, and 10 mM DTT) , the kinetic parameters were determined for biotinylated H4 (B-H4) (residues 1-24) by varying the peptide concentration (0–1.75 μM) and keeping the AdoMet at a saturation concentration of 10 μM. Apparent Km values were determined for AdoMet in reactions with 2 μM of B-H4 by varying the AdoMet concentration (up to 12 μM). The reactions were started by the addition of a mixture of $^{3}$H-AdoMet and unlabeled AdoMet. The reaction mixtures were incubated for 20 min at 23 °C and quenched by adding 10 μl of 7.5 M guanidinium hydrochloride. 10 μl of quenched reaction mixture was spotted onto streptavidin-coated membrane squares (SAM2® Biotin capture membrane, Promega). The membrane was washed three times in 2 M NaCl for 2 min each time, then in water three times for 30 s each. The membrane was dried at 50 °C for 1 h and then each spotted square was cut from the membrane and placed in a scintillation vial. The amount of methylated peptide was quantified by tracing the radioactivity (cpm) as counted by a TriCarb liquid scintillation counter (PerkinElmer Life Sciences).

**5. $IC_{50}$ Determination of (*R*)-1 against PRMT6-WT and -mutants**

$IC_{50}$ values of (*R*)-**1** (data summarized in Table S6) were determined using the radiometric method.[7] The pre-incubation time was 20 min.

**6. Biochemical assays**

$IC_{50}$ values inhibiting PRMT6 activity were also determined using the radiometric method.[8] The pre-incubation time was 2 h for all biochemical assays except the ones for Table S6 as noted above.

**7. Mass Spectrometry Analysis for Assessing the covalent binding:**

PRMT6 was incubated with 20 molar excess of the compound SGC6870 for 1hr at RT, followed by adding 0.1% trifluoroacetic acid (aq.). The samples were separated over a HPLC column over a 5-95% acetonitrile/water gradient and analyzed using an Agilent LC/MSD Time-of-Flight Mass Spectrometer equipped with an electrospray ionization source.

**8. Selectivity assays**

The effect of compounds on 33 methyltransferase (MT) activities were assessed as previously described.[9] Selectivity assay of 44 non-epigenetic targets (kinases, GPCRs, ion channels, and transporters) were conducted by Eurofins (https://www.eurofinsdiscoveryservices.com/cms/cms-content/services/adme-tox).

**9. Cellular PRMT6 assay**

HEK293T cells were grown in 12-well plates (2e5cells/well) in DMEM supplemented with 10% FBS, penicillin (100 units/ml) and streptomycin (100 μg/ml). Next day cells were transfected with FLAG-tagged PRMT6/mutant V86K, D88A PRMT6 (1 μg of DNA per well) using jetPRIME® transfection reagent (Polyplus-Transfection), following manufacturer instructions. After 4 h media were removed, and cells were treated with compounds. After 20 h, media was removed, and cells were lysed in 100 μl of lysis buffer (in mM: 20 Tris-HCl pH=8, 150 NaCl, 1 EDTA, 10 $MgCl_2$, 0.5% Triton-X100, 12.5 U/ml benzonase (Sigma), complete EDTA-free protease inhibitor cocktail (Roche). After 1 min. incubation at RT, SDS was added to the final 1% concentration. Total cell lysates were resolved in 4-12% Bis-Tris Protein Gels (Invitrogen) with MOPS buffer (Invitrogen) and transferred in for 1.5h (80 V) onto PVDF membrane (Millipore) in Tris-Glycine transfer buffer containing 20% MeOH and 0.05% SDS. Blots were blocked for 1h in blocking buffer (5% milk in 0.1% Tween 20 PBS) and incubated with primary antibodies: mouse anti-H4 (1:1000, Abcam #174628), rabbit anti-H4R3me2a (1:1000 Active Motif #39705), mouse anti-FLAG (1:5000, Sigma #F1804), mouse anti-H3 (1:1000, Abcam #174628), rabbit anti-H3R2me2a (1:1000, Millipore #04-808) in blocking buffer o/n at 4ºC. After five washes with 0.1% Tween 20 PBS the blots were incubated with goat-anti rabbit (IR800 conjugated, LiCor #926-32211) and donkey anti-mouse (IR 680,

LiCor #926-68072) antibodies (1:5000) in Odyssey Blocking Buffer (LiCor) for 1h at RT and washed five times with 0.1% Tween 20 PBS. The signal was read on an Odyssey scanner (LiCor) at 800 nm and 700 nm.

**10. Cell viability assay**

1 x $10^4$ cells (HEK293T, PNT2 and MCF7) were seeded on 96-well plates. After 24 h, the serial diluted compounds (SGC6870 and SGC6870N) from 10 μM were treated for 3 days. Cell viability was evaluated by CCK-8 (Cell counting kit-8, WST-8). Briefly, 10 μL/well of CCK-8 was treated and then incubated for 4 h at 37 °C. The absorbance was recorded by Infinite F PLEX plate reader (TECAN, Morrisville, NC, USA) at 450 nm. GraphPad Prism 8 has been used for data analysis and experimental results were shown as the average ± SD from three replicate experiments.
